# First report of ‘*Candidatus* Liberibacter asiaticus’ affecting sour orange in urban areas of Mayabeque, Cuba

**DOI:** 10.1101/2020.03.25.008623

**Authors:** Edel Pérez-López, Tim J. Dumonceaux

## Abstract

‘*Candidatus* Liberibacter asiaticus’ (CLas) is an unculturable, Gram-negative, phloem restricted plant pathogenic bacterium associated with a very serious disease of citrus worldwide known as Citrus Huanglongbing (HLB). CLas is widely spread in the Americas. In Cuba, CLas has been associated with HLB symptoms and has seriously affected the Cuban citrus industry. In this short communication we discuss the identification of CLas-infected sour orange in urban areas of Mayabeque Province in Cuba, an area previously unexplored for the presence of HLB, and a host widely cultivated in gardens and yards along Cuba. We used for the first time the bacteria molecular barcode chaperonin-60 universal target (*cpn60* UT) to identify and to detect CLas in HLB-symptomatic host plants.

---

‘*Candidatus* Liberibacter asiaticus’ (CLas) is a Gram-negative organism and one of the species within the genus ‘*Candidatus* Liberibacter’ in the alpha subdivision of the Proteobacteria (Jagoueix et al. 1994). Along with the species ‘*Candidatus* Liberibacter americanus’ (CLam) and ‘*Candidatus* Liberibacter africanus’ (CLaf), CLas has been associated with the disease Citrus huanglongbing (HLB), a devastating disease considered to be one of the most damaging plant diseases worldwide, which is associated with substantial economic losses in Asia, Africa and the Americas (Farnsworth et al. 2014; Bové 2014, da Graça et al. 2015). CLas is widespread in all HLB-affected countries in the Americas, with reports in the United States, Mexico, Brazil, and Cuba, among others (Teixeira et al. 2005; Bové 2006; Martínez et al. 2007).

Besides the 16S rRNA operon, which is present in three copies in the ‘*Ca*. L. asiaticus’ (CLas) genome (Duan et al. 2009), diversity studies of CLas have been restricted to the 16S-23S rRNA intergenic spacer region, the *omp* gene, the *β*-operon sequence, or bacteriophage-type DNA polymerase region (DNA pol) (Subandiyah et al. 2000; Bastianel et al. 2005; Ding et al. 2009; Furuya et al. 2010). The *cpn60* universal target (*cpn60* UT) (Goh et al. 1996), a fragment of approximately 550 bp, has been extensively used in the study of microbial communities, in the development of molecular diagnostic methods, and has been demonstrated to be a suitable molecular barcode for the domain Bacteria (Links et al. 2012; Pérez-López et al. 2016b). Partial *cpn60* gene sequences (500 to 550 bp) have been also useful to identify new species, and recently identified as an additional marker to improve the differentiation of plant pathogens within the genera ‘*Candidatus* Phytoplasma’ (Pérez-López et al. 2016a) and *Xanthomonas* spp. (Tian et al. 2016). Although CLas, CLam, and CLaf are the main pathogens associated with HLB disease (Bové 2014). The detection of sour orange (*Citrus aurantium*) trees showing HLB-related symptoms in urban areas in Mayabeque, Cuba along our previous successful experience identifying plant pathogens using the molecular marker *cpn60* UT motivated the development of specific primers targeting the CLas-*cpn60* UT sequence to confirm if the symptomatic plants were indeed affected by CLas.

Four sour orange plants (each plant representing a sample) showing diffuse chlorosis and corky veins, symptoms associated with HLB disease, were collected in Güines, Mayabeque, Cuba (22° 50′ 51″ N and 82° 01′ 25″ W) in May 2016, along with leaf tissue from one asymptomatic plant. Total DNA was extracted from the five plants as previously described (Pérez-López et al. 2016b), and used as template in PCR assays designed to amplify the *cpn60* UT sequence from CLas, a species known to be associated with HLB disease in Cuba (Table 1). Primers for CLas-*cpn60* UT PCR amplification were based on the *cpn60* UT primer annealing sites (Hill et al. 2004), but were adapted to ‘*Candidatus* Liberibacter’ sequences using full-length *cpn60* genes available in public databases (www.cpndb.ca and www.ncbi.nlm.nih.gov). Primers were developed using the interactive online tool Primer3web version 4.0 (http://primer3.ut.ee). Amplicons were purified using QIAquick® PCR Purification Kit (QIAGEN, Toronto, Canada) and directly sequenced using the amplification primers (Eurofins, CA). The product amplified from one of the samples was also ligated into the vector pGEM-T Easy (Promega, Madison, WI USA) following the manufacturer’s recommendations, transformed into chemically competent *E. coli* TOP10 (Invitrogen,ON, Canada), and sequenced using primers pair T7/SP6.

**Table 1.**
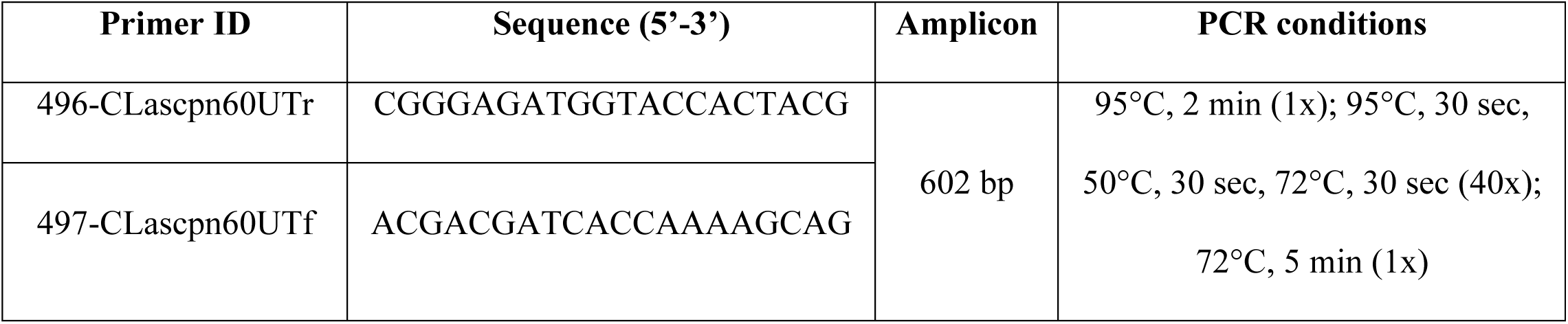
Primers and conditions used to amplify CLas-*cpn60* UT sequence.

Sequences generated from the four Cuban samples were assembled using the Staden package. Phylogenetic analysis of *cpn60* UT sequences (DNA and amino acid) of ClasCub samples, along with *cpn60* UT sequences from other ‘*Ca*. Liberibacter’ strains and other related bacteria were performed using maximum evolution method, using the close-neighbor-interchange (CNI) algorithm available in MEGA6 (Tamura et al. 2013), and bootstrapped 1000 times.

Mottle and corky veins were the main symptoms observed in sour orange collected in Mayabeque, Cuba (Fig. 1A). PCR targeting the amplification of the *cpn60* UT sequence of phytoplasmas was negative for symptomatic and asymptomatic samples (data not shown). Amplicons corresponding to the *cpn60* UT (∼600 bp) were amplified with 496-CLascpn60UTr/497-CLascpn60UTf primers from the four symptomatic citrus plants analyzed (CLasCub-1 to CLasCub-4) (Fig. 1B). Sequences generated were deposited to Genbank under the accession numbers KY581665 to KY581668 for CLasCub-1 to CLasCub-4, respectively. *cpn60* UT sequences obtained showed 100 % nucleotide identity among them, including the sequence cloned into the vector pGEM-T Easy (generated from LasCub-1), and 100 % identity with the CLas strains FL17 and psy62 from Florida (GenBank accession no. JWHA00000000 and CP001677, respectively), HHCA from California (GenBank accession no. JMIL00000000), the Chinese strains A4 and Guangxi-1 (GenBank accession no. JFGQ00000000 and CP004005, respectively), and the strain Ishi-1 from Japan (GenBank accession no. AP014595). Both phylogenetic trees (using amino acid and nucleotide sequences) showed distinction between the ‘*Ca*. Liberibacter’ species associated with HLB disease, and also a clear distinction from other related bacteria such as ‘*Ca*. L. solanacearum’ and *Naegleria gruberi* (Fig. 1C).

**Fig. 1.**
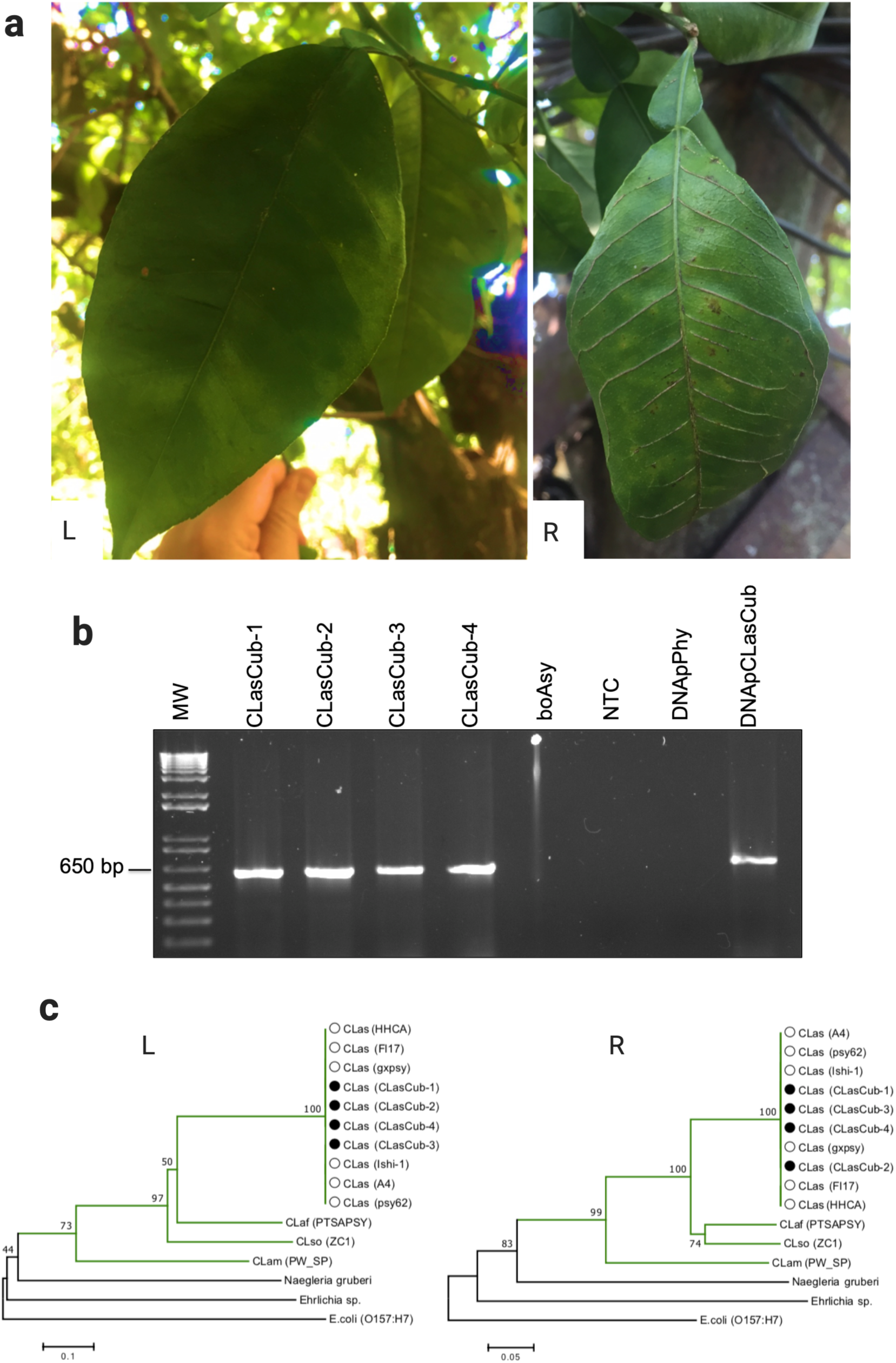
Identification of sour orange affected by ‘Candidatus Liberibacter asiaticus’ in Mayabeque, Cuba. (a) HLB-related symptoms observed in bitter oranges. Left (L): mottle observed in CLasCub-1. Right (R): corky veins observed in CLasCub-3. (b) Amplification of *cpn60* UT sequence using as template DNA extracted from four bitter orange trees from Cuba (CLasCub-1 to -4). Control used included bitter orange tree asymptomatic (boAsy), no template control (NTC), mixture of plasmid DNA containing *cpn60* UT of several phytoplasma groups and subgroups previously obtained by Dumonceaux et al. (2014) (DNApPhy), and plasmid DNA containing LasCub-*cpn60* UT (DNApCLasCub). Lane labelled MW represent Invitrogen 1 kb plus ladder. (c) Phylogenetic tree reconstructed through the neighbor-joining method of the *cpn60* UT nucleotide to the left (L) and amino acid to the right (R) sequences of ‘*Candidatus* Liberibacter’ strains identified in this study and previously characterized (GenBank accession number in the main text and Table 3). Other related bacterium included in the analysis was *Ehrlichia sp*. (KF977220), along the eukaryote *Naegleria gruberi* (XM_002680874). *E*.*coli* strain O157:H7 (NC_002655) was used as outgroup. Trees were bootstrapped 1000 times to achieve reliability. Bar, 5 substitutions in 100 positions (a), and 1substitution in 10 positions (b).

HLB disease was detected in Cuba in early 2005 and had serious negative consequences of citriculture (Martínez et al. 2009; Luis et al. 2009). All commercial varieties have been affected by the disease throughout the country, and the presence of HLB in urban areas has been reported before in Havana (Luis et al. 2009), but this is the first report in Mayabeque, a province that neighbours two main citrus-producing areas in Cuba: Ceiba and Jagüey Grande (Batista et al. 2014). Although there is not a national strategy, some companies and Citrus-producing areas opted to eradicate plantations affected followed by re-plantation (Batista et al. 2014), with a rigorous management based on the detection of new HLB-infected plants; for this reason, a strong surveillance is necessary.

Although protein-encoding genes are known to provide a better strain resolution compared to rRNA-encoding genes (Zeigler 2013), in CLas the close relationship among plant host (*Citrus* plants)-pathogen (CLas)-vector (*Diaphorina citri*) have influenced the high nucleotide identity among protein-encoding genes from different strains (Islam et al. 2012). Multilocus sequence analysis, microsatellite analysis, and genome sequencing are some of the strategies used to differentiate Las strains and even genetic groups among the strains reported worldwide (Islam et al. 2012; Katoh et al. 2014), but the high cost of those technologies is not affordable for countries such as Cuba. Specificity, sensitivity and reproducibility are important aspects that should be explored with a bigger set of samples in the near future.

The inclusion of the *cpn60* UT sequence as a marker to identify ‘*Candidatus* Liberibacter asiaticus’, and the use of the primers CLascpn60UTr/497-CLascpn60UTf designed in this study will improve the management of HLB in undeveloped and agricultural countries, allowing the detection of infected trees. To manage diseases associated with unculturable prokaryotes effectively, it is vital to understand the relationships between different species, emphasizing the importance of accurate microorganism identification.

## Acknowledgments

This work was supported by the Genomic Research and Development Initiative for the shared priority project on quarantine and invasive species. Edel Pérez-López thanks Agriculture and Agri-Food Canada Saskatoon Research Centre and the government of Canada for the time in Dumonceaux Lab.

